# Vasa vasorum lumen narrowing in brain vascular hyalinosis in systemic hypertension patients with ischemic stroke

**DOI:** 10.1101/2020.10.16.342741

**Authors:** Sergiy G. Gychka, Nataliia V. Shults, Sofia I. Nikolaienko, Lucia Marcocci, Nurefsan E. Sariipek, Vladyslava Rybka, Tatiana A. Malysheva, Vyacheslav A. Dibrova, Yuichiro J. Suzuki, Alexander S. Gavrish

## Abstract

Ischemic stroke is a major cause of death among patients with systemic hypertension. The narrowing of the lumen of the brain vasculature contributes to the increased incidence of stroke. While hyalinosis represents the major pathological lesions contributing to the vascular lumen narrowing and stroke, the pathogenic mechanism of brain vascular hyalinosis has not been well characterized. Thus, the present study examined the postmortem brain vasculature of human patients who died of ischemic stroke due to systemic hypertension. Hematoxylin and eosin staining and immunohistochemistry showed the occurrence of brain vascular hyalinosis with infiltrated plasma proteins along with the narrowing of vasa vasorum and oxidative stress. Transmission electron microscopy revealed the endothelial cell bulge protrusion into the vasa vasorum lumen and the occurrence of endocytosis in the vasa vasorum endothelium. The treatment of cultured microvascular endothelial cells with adrenaline also promoted the formation of the bulge as well as endocytic vesicles. siRNA knockdown of sortin nexin-9 (a mediator of clathrin-mediated endocytosis) inhibited the adrenaline-induced endothelial cell bulge formation. Adrenaline promoted protein-protein interactions between sortin nexin-9 and neural Wiskott–Aldrich Syndrome protein (a regulator of actin polymerization). We propose that endocytosis-depending endothelial cell bulge narrows the vasa vasorum, resulting in ischemic oxidative damage to the cerebral vessels, the formation of hyalinosis, the occurrence of ischemic stroke, and death in systemic hypertension patients.

## Introduction

Stroke is a leading cause of long-term disability and death worldwide. In the United States alone, a stroke occurs every 40 seconds and stroke-induced death occurs every four minutes. Every year, more than 795,000 Americans have a stroke, resulting in 140,000 deaths.^1–3^ By 2030, an additional 3.4 million U.S. adults are expected to have stroke (20.5% increase in prevalence from 2012).^4,5^ About 87% of all stroke incidences are ischemic stroke, which occur when a blood vessel supplying the brain is obstructed. Systemic hypertension is the most important risk factor for the development of ischemic stroke.^1,6^ High blood pressure promotes the alterations of the vascular wall and increases the incidence of ischemic stroke and subsequent neurodegeneration.^7–10^

One important lesion that contributes to the lumen narrowing of the brain vessels is vascular hyalinosis, which refers to the thickening of the vascular wall due to deposits of homogeneous hyaline materials.^11,12^ The pathogenesis of the formation of hyaline includes the infiltration of plasma proteins such as apolipoprotein E (ApoE), α2-macroglobulin, fibrinogen, and immunoglobulin G into the vascular wall.^13,14^ The accumulation of these plasma components alters all the structural components of the vessel wall through the formation of fibrinoid lesions and causes the hyalinization of the vessel wall.^15–17^

Thus, the reversal and/or prevention of brain vascular hyalinosis in systemic hypertension patients have the therapeutic potential to reduce the incidence of ischemic stroke. However, the mechanism of brain vascular hyalinosis is not well understood. While the involvement of oxidative stress in various pathological features of stroke has been documented,^18^ whether oxidative stress occurs in hyalinosis lesions is unknown. The present study performed detailed histological analyses of the hyalinosis lesions in the postmortem brain tissues of human patients who died of ischemic stroke. We found that the brain vascular hyalinosis in human systemic hypertension patients who died of ischemic stroke is associated with the narrowing of the vasa vasorum that supplies the blood to the cerebral vascular wall and the occurrence of oxidative stress in both the vessel walls as well as the brain tissues.

## Materials and Methods

### Autopsy brain tissues from patients

Postmortem brain tissues were collected from 28 patients with a history of systemic hypertension and who died of ischemic stroke and 28 patients who died of ischemic stroke without systemic hypertension. The tissues were taken from the perifocal zone of the ischemic infarct in region of the middle cerebral artery of the frontal lobe. Clinical studies were approved by the regional committee for medical research ethics in Kiev, Ukraine (ethical code: 81, 2016) and performed under the Helsinki Declaration of 1975 revised in 2013 or comparable ethical standards. The participants gave written informed consent.

### Experimental animals

Male and female spontaneously hypertensive stroke prone (SHRSP) rats were purchased from Charles River Laboratories International, Inc. (Wilmington, MA, USA). Animals were fed normal rat chow. The Georgetown University Animal Care and Use Committee approved all animal experiments, and the investigation conformed to the National Institutes of Health (NIH) Guide for the Care and Use of Laboratory Animals.

### Histological measurements

The sections were resected to be approximately 10 μm thick and immersed in 10% buffered formalin at room temperature. The fixed tissue samples were paraffin embedded, sectioned at 6 μm with a microtome, mounted on glass slides, and analyzed histochemically and immunohistochemically. Tissue sections were subjected to hematoxylin and eosin (H&E) staining to determine the general morphology of the brain vessels, Zerbino-Lukasevich staining to detect the presence of fibrin to assess the vascular lesions, and immunohistochemistry using the anti-ApoE antibody (Abcam, Cambridge, UK) and anti-malondialdehyde (MDA) antibody (Abcam). The vessels smaller than 25 μm or larger than 150 μm in external diameter were excluded from analysis.

### Transmission Electron Microscopy (TEM)

Brain tissues were collected from patients who underwent neurosurgery and immediately fixed in a solution containing 4% paraformaldehyde and 0.5% glutaraldehyde/0.2 M cacodylate. Samples were then post-fixed with 1% osmium tetroxide and embedded in EmBed812. Ultrathin sections were stained with uranyl acetate and lead citrate and examined in a Philips EM-400T Transmission Electron Microscope at 80 kV with the TIA software.

### Cell culture

Human microvascular endothelial cells were purchased from ScienCell Research Laboratories (Carlsbad, CA, USA) and were cultured in accordance with the manufacturer’s instructions in 5% CO_2_ at 37°C. Cells in passages 3-6 were used.

siRNA experiments were performed using the Santa Cruz Biotechnology (Dallas, TX, USA) system in accordance with the manufacture’s instructions. Briefly, cells grown on 6-well plates were incubated with 1 μg siRNA and siRNA Transfection Reagent in siRNA Transfection Medium for 5 hours. Equal volume of growth medium containing 2 times the normal serum, growth supplements and antibiotics were added and cells were grown for 2 days.

### Immunoprecipitation and Western blotting

Cell lysates were immunoprecipitated with the mouse polyclonal anti-sortin nexin 9 (SNX9) antibody (Santa Cruz) and SureBeads Protein G Magnetic Beads (Bio-Rad Laboratories, Hercules, CA, USA) for 1 h at room temperature. Immunoprecipitation using SureBeads was performed in accordance with the manufacturer’s instructions. Samples were electrophoresed through a reducing SDS polyacrylamide gel and electroblotted onto a nitrocellulose membrane. Membranes were blocked and incubated with the antibody to detect neural Wiskott-Aldrich Syndrome protein (N-WASp; MilliporeSigma, Burlington, MA, USA) using horseradish peroxidase-linked secondary antibodies and an Enhanced Chemiluminescence System (GE Healthcare Bio-Sciences, Pittsburgh, PA, USA). Autoradiography was performed using UltraCruz Autoradiography Films (Santa Cruz Biotechnology). The developed films were scanned and optical densities of protein bands were quantified using NIH ImageJ.

### Statistical analysis

Statistical analysis was performed with the IBM SPSS Statistics version 23.0 software. Significant differences between two groups were determined by the Independent Samples Mann-Whitney U test.

## Results

### Vasa vasorum narrowing in vascular hyalinosis lesions in the brain of systemic hypertension patients who died of ischemic stroke

Hyalinosis occurs in the brain vasculature in late-stage systemic hypertension preceding ischemic stroke. Hyalinosis is characterized by the death of vascular cells, ultimately leading to the infiltration and accumulation of plasma proteins, forming glass-like materials that occlude the blood vessel lumen and interfere with the gas exchange between the blood and brain tissues. The Zerbino-Lukasevich staining results shown in Fig. 1A demonstrate the fibrin infiltration into the vascular wall from the lumen during vascular wall hyalinization. The higher magnification image in Fig. 1A (right panel) shows the presence of fibrins in the blood plasma, the precipitation of fibrins at the luminal surface of blood vessels, and infiltration into the vessel wall. ApoE is another protein that infiltrates the hyalinosis lesion.^13^ Immunohistochemistry for ApoE revealed positive stains in the vascular wall in association with increased wall thickness due to high blood pressure in systemic hypertension patients who died of ischemic stroke (Fig. 1B).

**Figure 1:**
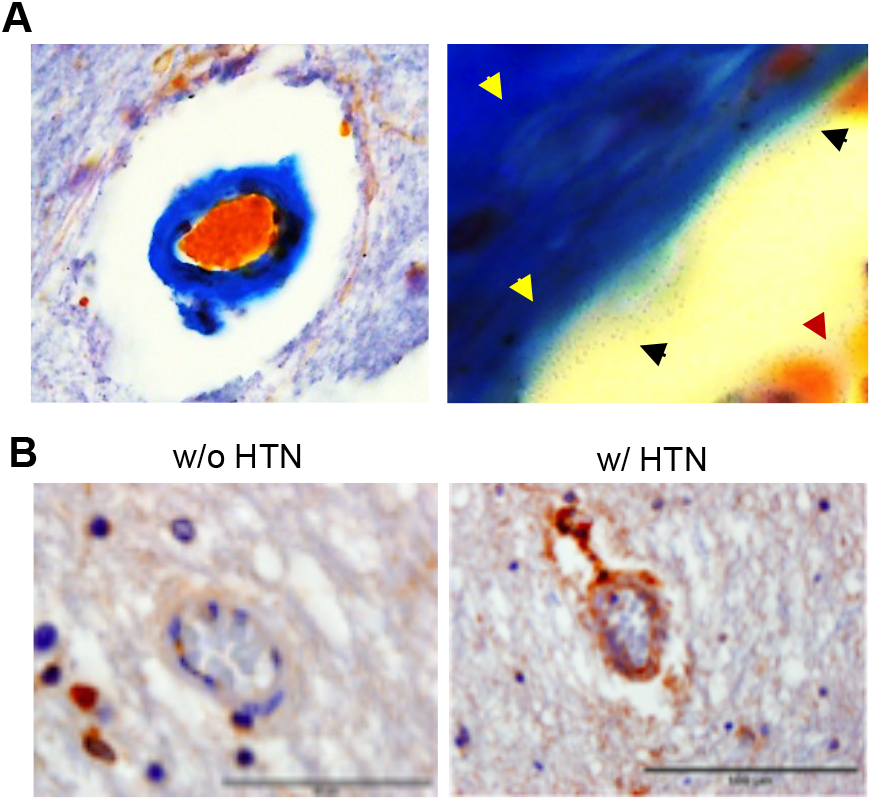
Vascular hyalinosis in systemic hypertension patients who died of ischemic stroke. (A) A representative Zerbino–Lukasevich staining of patients with systemic hypertension who died of ischemic stroke. The left panel shows the infiltration of plasma fibrin into vessel wall. In the right panel, higher magnification showing fibrin in the blood (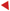), at the vessel surface (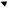), and in the vessel wall (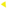). (B) Representative immunohistochemistry results using the ApoE antibody in patients who died by ischemic stroke without (w/o) or with (w/) hypertension (HTN). Positive ApoE stains in the intima, media and adventitia layer are found only in w/HTN. Magnifications x400. Representative images of N=8 patients per group.

In these systemic hypertension patients who died of ischemic stroke, H&E stain shows the hyalinosis lesion, with significant cerebral vessel wall thickening (Figs. 2A and 2B) and lumen narrowing (Figs. 2A and 2C). Histological analysis of the adventitia layer of the cerebral vessel revealed that patients who died of ischemic stroke in association with systemic hypertension exhibited significant structural changes of the vasa vasorum. Fig. 2A shows representative H&E stain images of vasa vasorum of cerebral vessels from patients who died of ischemic stroke due to systemic hypertension in comparison with patients without systemic hypertension. In patients with systemic hypertension, endothelial cells of the intima layer of the vasa vasorum were found to be enlarged and the vasa vasorum media layer is thickened. Morphometric analysis of the vasa vasorum (<25 μm) revealed the wall thickening (Fig. 2D) and a reduction in the lumen area (Fig. 2E) of the vasa vasorum in patients who died of ischemic stroke due to systemic hypertension.

**Figure 2:**
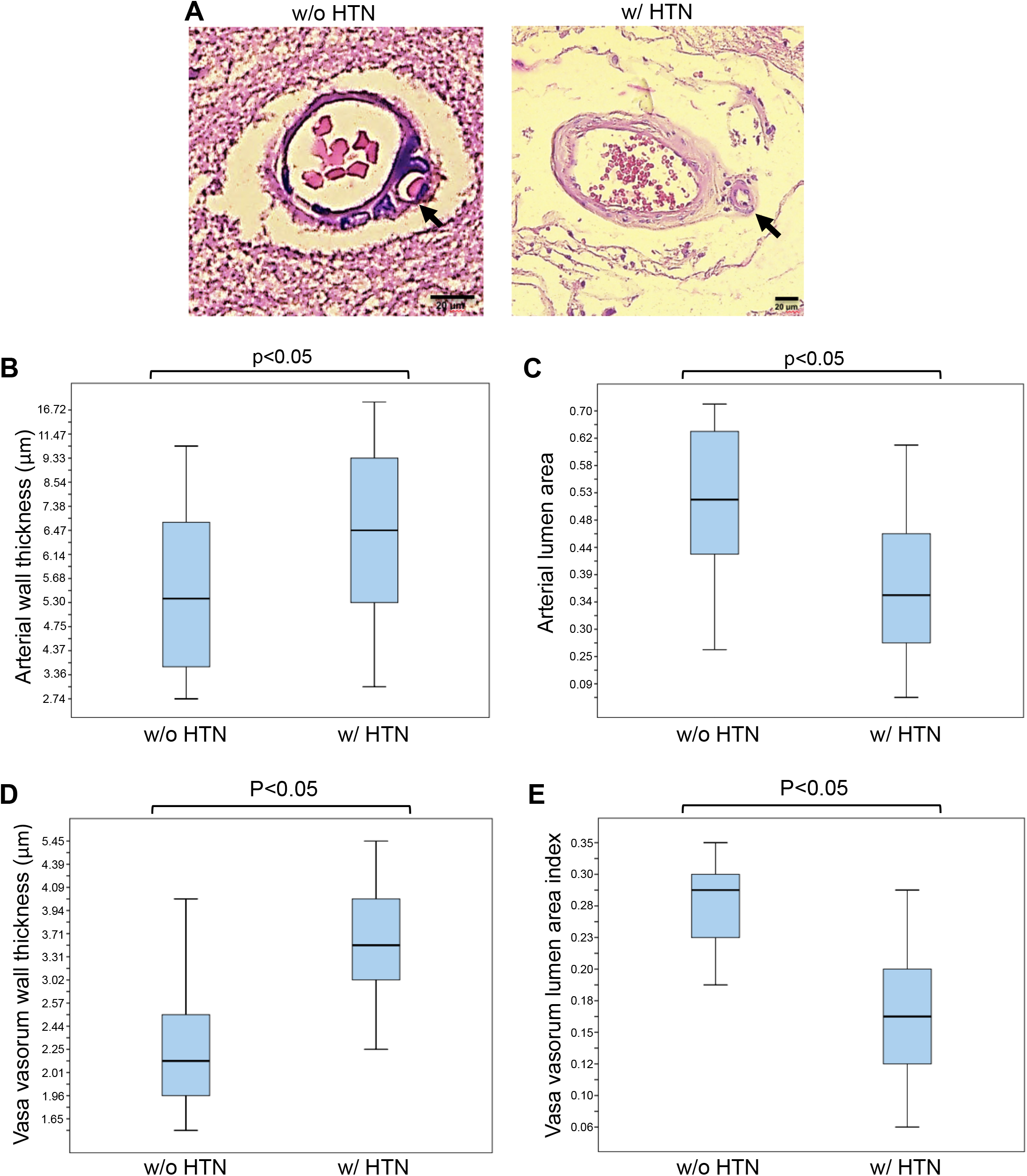
H&E staining of vasa vasorum in the brain of patients who died of ischemic stroke. (A) Ischemic stroke without (w/o) systemic hypertension and with (w/) systemic hypertension. Representative images of N=28 patients per group. (B) Quantification of the vessel wall thickness. (C) Quantification of cerebral vascular lumen area index that was calculated using the equation, internal surface area/external surface area. N=8 for without (w/o) HTN and N=6 for with (w/) HTN. (D) Quantification of the vasa vasorum wall thickness. (E) Quantification of vasa vasorum lumen area index that was calculated using the equation, internal surface area/external surface area. N=8. Mann-Whitney U Test indicated that values in the box plots are significantly different at p<0.05.

Since the vasa vasorum supplies the blood to the walls of cerebral blood vessels (>50 μm), the narrowing of the vasa vasorum may contribute to the initiation and/or progression of cerebral vessel hyalinosis by limiting the blood flow as well as oxygen delivery to the brain, promoting oxidative stress. Indeed, immunohistochemistry using the antibody against malondialdehyde (MDA), a lipid peroxidation product, showed that patients who died of ischemic stroke with systemic hypertension, but not without systemic hypertension, have positive MDA staining in both cerebral vessel walls and in the neuronal tissues (Fig. 3).

**Figure 3:**
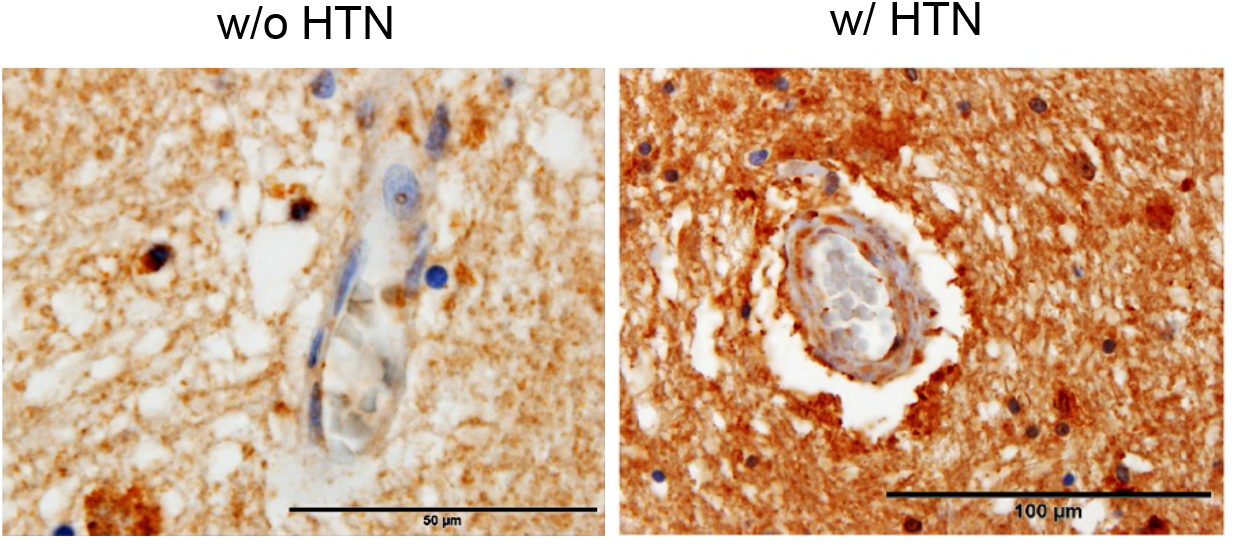
Immunohistochemistry for MDA. Ischemic stroke without systemic hypertension (w/o HTN) and with systemic hypertension (w/HTN). Magnifications ×400. Representative images of N=8 patients per group.

### Endothelial cell bulge protusion into vasa vasorum in the vascular hyalinosis lesions in the brain of systemic hypertension patients who died of ischemic stroke

Our detailed examinations of the hyalinosis lesion in brain tissues collected from human patients who died of ischemic stroke led us to realize that endothelial cell structural changes contribute to the occlusion of the vasa vasorum due to the formation of the “bulge” structure that protrudes from endothelial cells into the lumen of the vasa vasorum (Fig. 2A). This lead us to formulate a novel hypothesis that the occlusion of this small vessel network supplying the blood to cerebral vascular wall tissues contributes to the death of vascular cells and the formation of hyalinosis. This endothelial cell bulge structure was also visualized by TEM in the vasa vasorum in the hyalinosis lesion of brain vessels of systemic hypertension patients who underwent neurosurgery to remove the hematoma to treat hemorrhagic stroke (Fig. 4A). In the vasa vasorum endothelial cells of brain vasculatures of these patients, we also noted the formation of endocytic vesicles (Fig. 4B).

**Figure 4.**
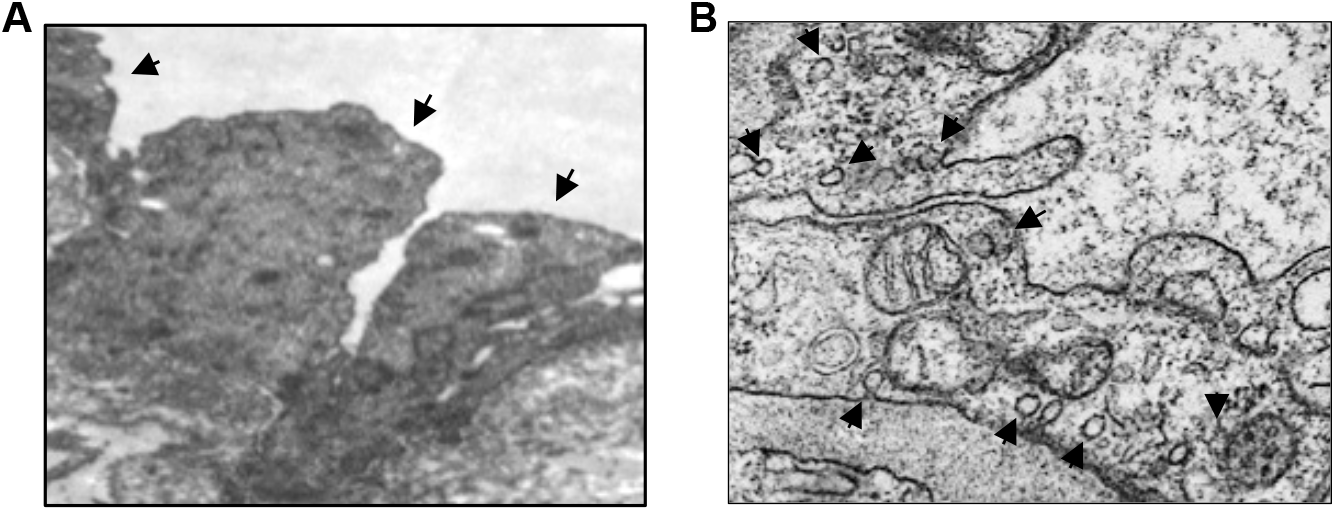
TEM images of vasa vasorum of hyalinosis lesion of brain vessels of systemic HTN patients. (A) Arrows indicate the presence of endothelial cell bulge. (B) Arrows indicate the presence of endocytotic vesicles. Representative images of N=6 patients.

Also, in cultured human microvascular endothelial cells, the adrenaline treatment promoted the formation of endocytic vesicles as visualized by TEM (Fig. 5A) as well as the bulge-like structure as observed via light microscopy (Fig. 5B). Immunofluorescence staining showed that the bulge formation induced by adrenaline is associated with the reorganization of actin filaments (Fig. 5C). We hypothesized that adrenaline activated the clathrin-mediated endocytosis that, in turn, elicits protein-protein interactions between SNX9 (a mediator of endocytosis) and N-WASp (a regulator of actin polymerization) (Yarar, 2007; Pollitt, 2009; Lundmark, 2009), resulting in actin filament structural changes in endothelial cells. In support of this, we found that siRNA knockdown of SNX9 inhibited the adrenaline-induced bulge formation (Fig. 5B) and that adrenaline promoted SNX9/N-WASp interactions (Fig. 5D).

**Figure 5:**
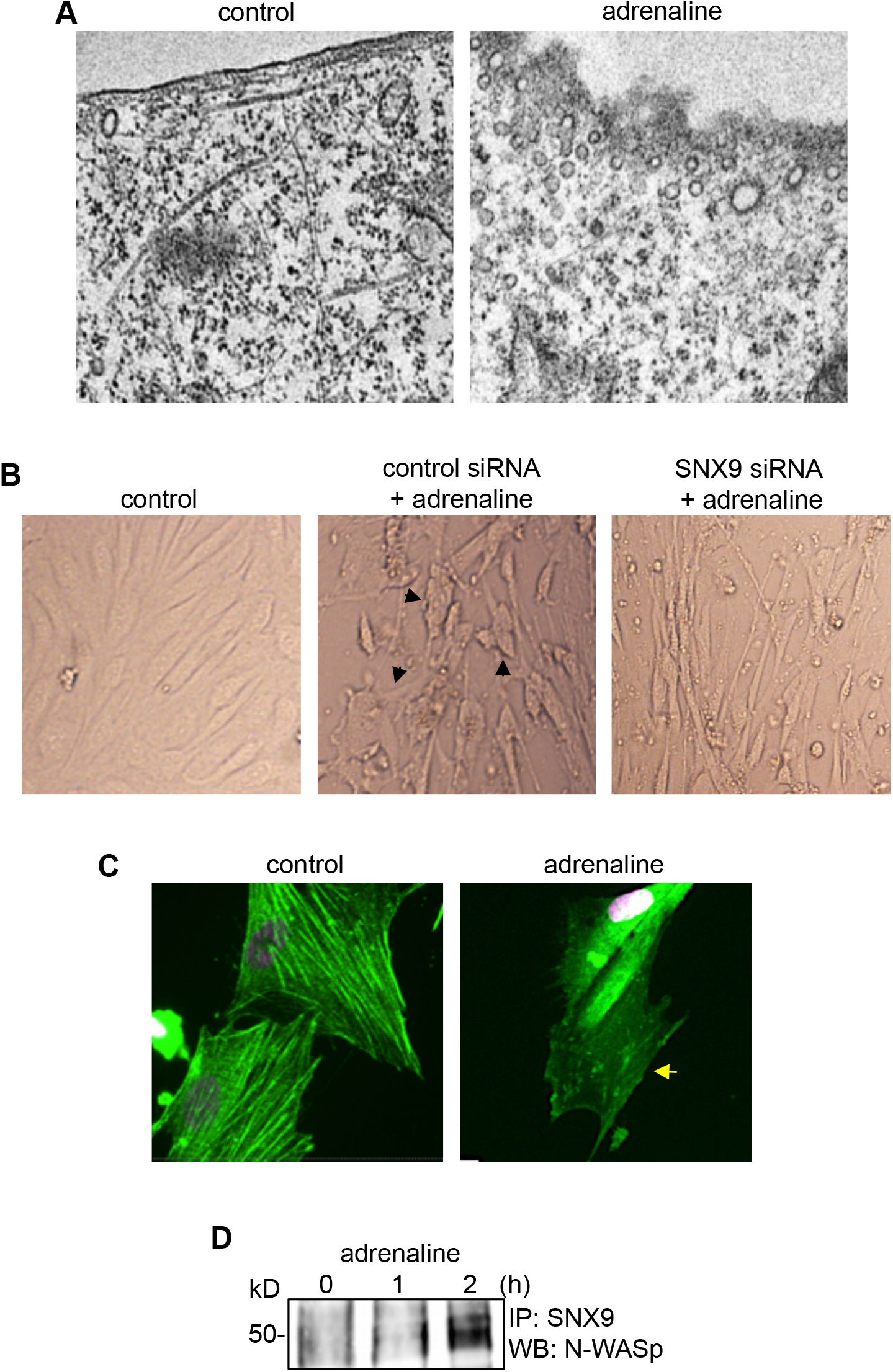
Studies of cultured human microvascular endothelial cells. (A) Transmission electron microscopy images of control cells and cells exhibiting adrenaline-induced formation of endocytotic vesicles. Cells were treated with adrenaline at 10 μM for 4 h. (B) siRNA knockdown of SNX9 inhibits the adrenaline-induced bulge formation in human microvascular endothelial cells. Cells were treated with siRNA for 2 days and then treated with adrenaline at 10 μM for 4 h. (C) Immunofluorescence staining of actin in untreated control and endothelial cells treated with adrenaline (10 μM; 4 h). Arrow indicates reorganization of actin filaments. (D) Adrenaline promotes SNX9/N-WASp interactions. Human microvascular endothelial cells were treated with 10 μM adrenaline for 0, 1 or 2 h. Cell lysates were subjected to immunoprecipitation (IP) with mouse SNX9 IgG followed by Western blotting (WB) with rabbit N-WASp IgG.

SHRSP rats exhibited the lesions that are indicative of brain vascular hyalinosis at 8 weeks of age (Fig. 6A). In TEM images of the brain vessels of these rats, we also observed the endothelial cell protrusion into the lumen of vasa vasorum (Fig. 6B) as well as endocytosis in the vasa vasorum endothelial cells (Fig. 6C).

**Figure 6:**
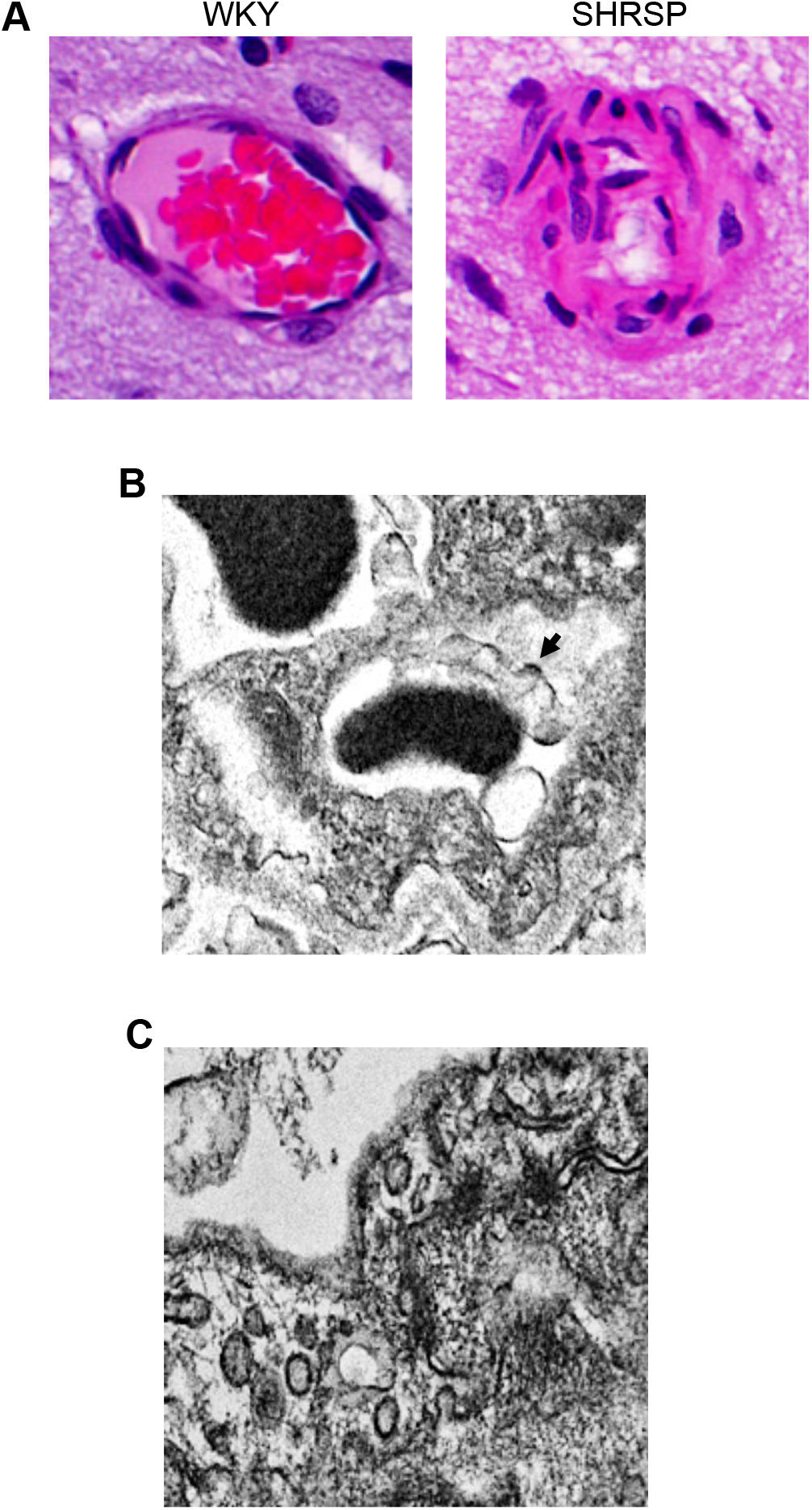
Spontaneously Hypertensive Stroke Prone (SHRSP) rats. (A) Representative H&E staining images of brain vessels in Wistar-Kyoto (WKY) control and SHRSP rats at 8 weeks of age. (B) A representative TEM image of endothelial cell bulge protrusion into the lumen of vasa vasorum in SHRSP rats. (C) A representative TEM image of endocytosis in the endothelium of vasa vasorum of SHRSP rats. N=4 rats.

## Discussion

In postmortem brain tissues obtained from patients who died of ischemic stroke due to systemic hypertension, we performed histological analyses to define the occurrence of brain vascular hyalinosis. In such lesions, plasma proteins are deposited in the vessel wall and the indication of oxidative stress was detected. We also found that the lumen of the vasa vasorum that supplies the blood to the wall of cerebral vessels was narrowed.

According to our result, lipid peroxidation products reside in both the hyalinosis lesions in the vessels as well as in the brain tissues. Immunohistochemistry using the MDA antibody could stain free MDA or MDA-protein adducts. Thus, it is not yet clear if lipid peroxidation actually occurs in the membrane systems associated with the hyalinosis lesions or the MDA-bound protein migrates to the hyalinosis lesions. Plasma proteins that are infiltrated may be oxidized and form oxidation-dependent aggregates, which may then be accumulated as hyaline.

The major finding of this study is that in addition to the cerebral arterial vessels, vasa vasorum is also narrowed. These results may suggest that ischemia-reperfusion injury due to the narrowing of vasa vasorum trigger oxidative stress damage to the cerebral vascular walls, resulting in the subsequent occlusion of the cerebral arteries and ischemic insult to the brain tissue.

The present study provide evidence to support the mechanistic hypothesis as depicted in Fig. 7 that the endothelial cell cytoskeletal rearrangement forms the bulge-like structure in the lumen of the vasa vasorum through adrenaline-induced endocytosis and SNX/N-WASp interaction-mediated actin reassembly, contributing to the occlusion of these small vessels that supply oxygen to the cerebral vascular wall.

**Figure 7:**
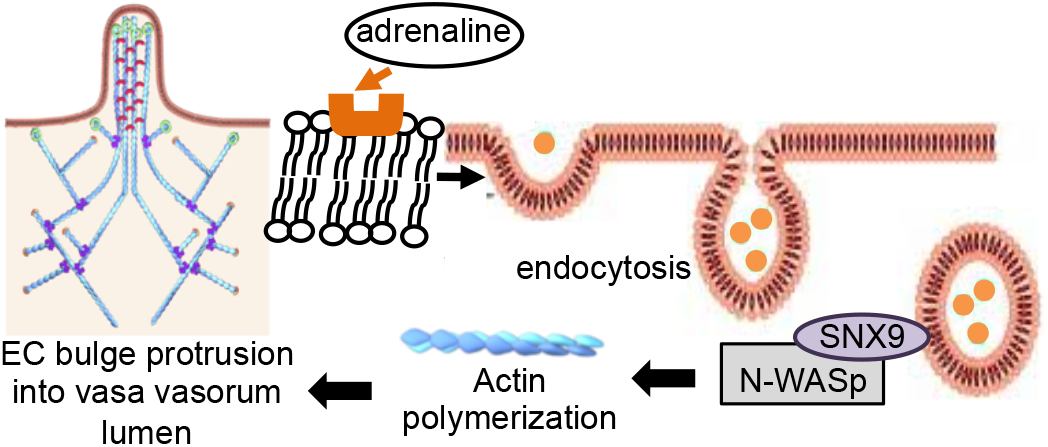
Scheme depicting the proposed mechanism of endothelial cell (EC) bulge protrusion into vasa vasorum lumen. In systemic hypertension patients, adrenaline promotes endocytosis-dependent interactions of SNX9 and N-WASp that activate actin polymerization and EC bulge protrusion into the lumen of vasa vasorum, resulting of the narrowing of the lumen of the vasa vasorum.

## Conclusion

In summary, the present study provided histological characterizations of human brain vascular hyalinosis in systemic hypertension patients who died of ischemic stroke. We propose that the vasa vasorum narrowing results in oxidative stress to the cerebral vessels, contributing to the formation of brain vascular hyalinosis, the occurrence of ischemic stroke, further oxidative stress to the neuronal cells, and eventual death in systemic hypertension patients. Further understanding this pathological lesion is critical to developing new and effective therapeutic strategies to reduce the risk of ischemic stroke in systemic hypertension patients, which comprise a large population worldwide.

## Funding

This work was supported in part by NIH (R01HL072844, R21AI142649, R03AG059554, and R03AA026516) to Y.J.S. The content is solely the responsibility of the authors and does not necessarily represent the official views of the NIH.

## Conflicts of Interest

The authors declare no conflict of interest.

## Notes

### Competing Interest Statement

The authors have declared no competing interest.

